# Cortical Tension Links Curvature to Tissue Growth in the Cellular Potts Model

**DOI:** 10.1101/2025.05.26.656125

**Authors:** Kai Lennard Fastabend, Cécile M. Bidan, John W. C. Dunlop, Philip Kollmannsberger

**Affiliations:** Biomedical Physics, Heinrich Heine University Düsseldorf, 40225 Düsseldorf, Germany; Max Planck Institute of Colloids and Interfaces, Dept. of Biomaterials, 14476 Potsdam-Golm, Germany; MorphoPhysics Group, Department of the Chemistry and Physics of Materials, Paris-Lodron University of Salzburg, 5020, Salzburg, Austria

**Keywords:** Cellular Potts Model, tissue surface tension, biomechanical growth factors

## Abstract

The growth of biological tissue is sensitive to the physical properties of the environment. For example, the growth rate of contractile tissue under geometric confinement is proportional to local curvature as a result of tissue surface tension. It is not known how local cell behavior is coordinated to give rise to such behavior. Here, we use computer simulations based on the Cellular Potts Model (CPM) to investigate the role of cell adhesion and cortical contractility for curvature-dependent tissue growth. Our results show that tension-dependent local proliferation leads to the observed macroscopic curvature-driven growth kinetics via tissue surface tension, independent of soluble growth factors.

## Introduction

Growth and regeneration of biological tissues are complex and poorly understood phenomena. Cells migrate, divide, and deposit matrix in the available space, but little is known about how the coordination of cell behavior in space and time leads to higher-order organization. The activity of a single cell and the chemical and mechanical signals it sends and receives depend on the microenvironment defined by other cells, giving rise to a reciprocal feedback loop (1, 2) in which early tissue serves as substrate for the growth of later tissue. A better quantitative understanding of how the chemical, mechanical and geometric properties of the substrate determine tissue organization would greatly benefit the design of engineered scaffolds to guide the formation of functional tissue. A particularly interesting example of collective control of cellular dynamics is the curvature-dependent growth of tissue under geometric confinement. Contractile adherent cells such as fibroblasts (skin cells) or osteoblasts (bone cells) form a dense tissue in concave pores or clefts with a growth rate strictly proportional to the local curvature. This surprising relationship was first observed in an *in vitro* model of osteoid deposition by mouse preosteoblasts on bone-like substrates and has since been reproduced in different systems using a variety of cell types (4–9). After an initial lag phase that depends on pore size and substrate material, the rate of growth is the same for a given cell type and pore geometry (10). A simple geometric construction that describes cells as linear tensile chords growing layer by layer explains the relationship between curvature and the movement of the interface (11), suggesting surface tension as the mechanism behind curvature-driven tissue growth. This tissue surface tension is a result of the interplay of cell contractility and pore geometry, as confirmed by experiments (6, 8, 12). Surface tension and curvature together lead to an outward-directed Young-Laplace force, driving the evolution of the tissue toward a minimal surface (9, 13). On a macroscopic level, surface tension explains not only growth kinetics but also cell and matrix alignment (14), in excellent agreement with experimental observations (15, 16). Tissue growth under geometric confinement is thus a example of the role of curvature in biological processes (17).

However, it is not clear how macroscopic curvature guides cellular decision-making toward proliferation and matrix production. Generally, adherent cells migrate, divide, differentiate and deposit matrix fibers in response to chemical and mechanical signals. On flat substrates, cell adhesion and spreading are not only required for proliferation (18), but higher substrate stiffness also leads to increased cytoskeletal tension and promotes differentiation into a more contractile phenotype (19, 20). The actomyosin cell cortex is an important regulator for cell shape and tissue patterning (21, 22), and the shape of invidual cells confined to adherent micropatterns as well as the shape of microtissues confined by cantilevers can be explained by cortical contractility (23, 24). During curvature-driven tissue growth, a gradient of cell proliferation, myofibroblast differentiation and matrix phenotype emerges (8, 25), suggesting a key role of cortical contractility not only for the macroscopic surface tension and curvature dependence, but for mediating macroscopic curvature down to the level of individual cells and their behavior, thus closing the feedback loop of dynamic reciprocity.

This hypothesis can be tested using cell-level computer simulations by modulating how mechanical signals control cell behavior and observing the outcome, which can not easily be done in experiments. Depending on the specific question, different approaches to model tissues with cellular resolution exist. For example, vertex-based models are well suited to study epithelial tissues with straight cell edges (26). By explicitly including cortical tension in the energy term, vertex models provide valuable insights into epithelial topology (27), but are less suitable for highly dynamic growing tissues and irregular cell shapes. Another class of models describes cells as simplified spheres or point-like objects without accounting for cellular morphology (28). Such models are, e.g., used to study spatial interactions and tissue topology during embryonal development (29) but do not capture the interplay with external geometric features.

A well-established cell-level computational modeling framework is the Cellular Potts Model (CPM) or Glazier-Graner-Hogeweg (GGH) model. Since its introduction in 1992 (30), it has led to important insights into how cell behavior influences the growth and patterning of complex biological tissues, reviewed in (31). Interactions of cells with other cells and the substrate are modeled by interfacial energies, while mechanical properties and contractility are included via energetic constraints. A lattice-based implementation allows for arbitrary cell shapes and a numerically efficient minimization of system energy under fluctuations. The CPM can, e.g., explain the role of cell adhesion and cortical contractility for eye patterning in drosophila (32) and the shape of single cells on micropatterns (33), making it a promising approach to investigate a possible link between curvature-driven tissue growth and cellular contractility.

In this study, we amend the CPM with a cortical tension-dependent cell proliferation rule to investigate the interplay of cell behavior, macroscopic scaffold curvature, and tissue growth. The model explicitly includes cell-cell and cell substrate adhesion, cortical contractility, and elasticity as separate terms in the system energy. The proliferation rule prohibits growth when cells are under compression, which is experimentally well established. The kinetics of biological growth is separated from the simulation timescale by averaging over fluctuations. We quantitatively compare the results of our simulations to experimental data of tissue growth *in vitro* and systematically explore which parameter ranges give rise to the observed behavior. All assumptions of the model are on the cellular level and are experimentally established (34), while the simulation results are compared to experiments on the tissue scale. Our results agree well with observations, suggesting a mechanobiological feedback loop involving cell adhesion, contractility and proliferation as mechanism behind curvature-driven tissue growth under geometric confinement.

## Materials and Methods

### Introduction to the Cellular Potts Model

The Cellular Potts Model (CPM), also referred to as Glazier-Graner-Hogeweg model (GGH) (31), is a cell-level modeling approach on a lattice, where sites with the same cell index form cells of variable shapes. Historically, the model is derived from the statistical mechanics of magnetic domains in the Ising and Potts models, with interaction energies between lattice sites of unequal spin values, or, in the case of the CPM, cell indices (30, 31). The dynamics of the simulation result from the minimization of the system energy in a stochastic Monte Carlo process that is based on pixel copy attempts defined by a Metropolis Algorithm. This results in a quasi-deterministic development of the system towards the energy minimum, defined by the energy function:

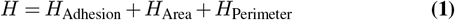

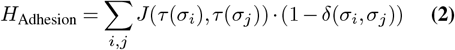

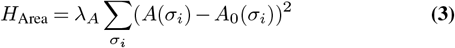

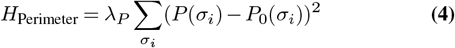

The energy function consists of different terms for adhesion, area, and perimeter constrains, which in their interplay determine cell shape and dynamics. The adhesion term sums up the interaction energies *J* of all lattice sites with neighboring sites of different cell indices. The contact energy *J* depends on the cell types *τ* of the contacting cells and defines whether it is an attractive or repulsive contact interaction. Increasing the size of the neighborhood determines the performance of the simulation and influences possible grid artifacts.

The area and perimeter constraints quantify an energy cost for the deviation of the area and perimeter of each cell from the target values *A*_0_ and *P*_0_, with an elastic potential that is scaled by the prefactors *λ*_*A*_ and *λ*_*P*_. Such an elastic area constraint was first introduced for the modeling of foams (35). In the context of a two-dimensional cell model, the area constraint acts like a surface tension that denotes an energy cost to changes of the area, while the perimeter constraint introduces a line tension to the contour of the cell. The force balance between surface tension and line tension determines the resulting cell shape. An example would be the formation of a 2D sphere due to the balance between surface tension *σ* and line tension *λ* where the radius is given by 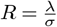.This is the 2D analog to the 3D Young-Laplace law, where the line tension is substituted by a surface tension, and the surface tension is exchanged by the internal pressure (33). In the biomechanical context of cell and tissue simulations, the area constraint introduces a resistance against compression (or expansion), while the elastic line tension determines the cell shape due to the restriction of the contour length. The perimeter constraint therefore acts like the cortical tension of the cell that results from the contractile actin-myosin network in the plasma membrane (22).

The quasi-deterministic development of the system down the gradient of energy is defined by pixel copy attempts in a Metropolis algorithm. For each pixel copy attempt, the value of a random lattice site is copied onto a random neighbor. If the new configuration after the pixel copy is energetically favorable, then the pixel copy attempt is accepted. If the new configuration increases the global energy of the system, the copy attempt is accepted with a probability that depends on the energy cost:

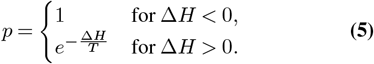

The result of this acceptance function is that the state of the system will diffuse down the gradient of energy with some statistical fluctuations that are scaled by the temperature *T*. Following the directions of possible pixel copy attempts along the cell boundary, one can use the local energy gradients for the calculation of forces to visualize the underlying mechanics of the simulation more intuitively (36). Fig. 1 visualizes conceptual equilibrium states of a suspended and adherent cell. In the context of cell simulations, the parameter *T* is typically referred to as the membrane fluctuation level. For each Monte Carlo step (MCS) of the simulation, multiple pixel copy attempts are performed sequentially. The number of pixel copy attempts per MCS is given by the number of pixels in the simulation box.

**Fig. 1.**
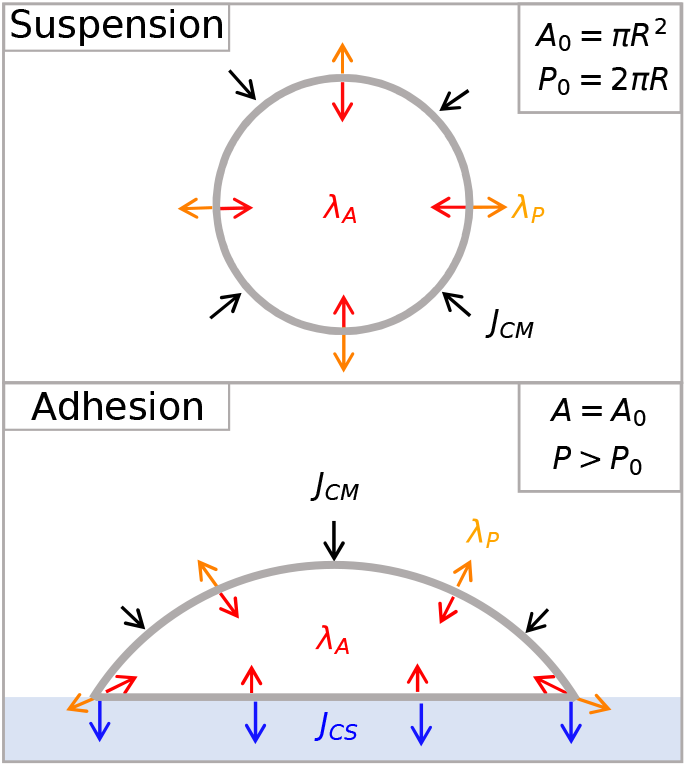
Conceptual visualization of suspended and adherent cells in force equilibrium under area and perimeter constraints and adhesion. The area constraint introduces a tension corresponding to the compressibility of the cell, while the perimeter constraint describes a line tension along its contour. The overall cell shape in suspension results from the force balance between area and line tension (top). Adhesion energies act as additional differential line tensions for specific interfaces and chance the force balance and resulting cell shape (bottom): substrate adhesion induces a change in cell shape similar to the wetting of a droplet on a hydrophilic substrate, since the cell-medium interface is energetically more costly than the cell-substrate interface. The resulting increase in perimeter leads to an increase in cortical tension, which is a biological signal required for proliferation of adherent cells. As described by Rens et al. (36), the indicated force directions reflect the gradients of the different terms of the energy function at the cell boundary.

The CPM is a powerful framework for the definition of biological models for tissues on an intermediate scale that consider single cells. To describe the behavior of growing tissues under geometric confinement, the model is easily expandable to contain cell proliferation and mitosis.

### Tension-Dependent Proliferation

Depending on the energy terms for cell elasticity and cellular adhesion, different cell shapes emerge during stochastic energy minimization. Assuming a single cell without adhesion, a round shape with radius *R* can be parameterized in continuous space by the target area *A*_0_ = *πR*^2^ and the target perimeter *P*_0_ = 2*πR*. Due to discretization errors of a continuous circle onto a square lattice, the perimeter *P*_0_ has to be rescaled by the prefactor *ε*_*P*_. Note that the value of the prefactor, which is defined by the quotient of discrete and continuous perimeter (*ε*_*P*_ = *P*_discrete_*/P*_continuous_) has a constant value of 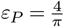 for all cell sizes (see supplement Fig. S1 for more detailed information on the discretization of circles). Using the formula for continuous circles and the previously introduced prefactor *ε*_*P*_, the relation between target area *A*_0_ and target perimeter *P*_0_ becomes:

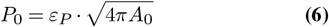

Apart from membrane fluctuations, a suspended cell (without medium interaction: *J*_CM_ = 0) will maintain its round shape, as this represents the most energetically favorable state where both perimeter and area requirements are satisfied. This situation changes when contact energies with the substrate or medium come into play, leading to alterations in cell shape. In the case of energetically favorable adhesion (negative *J*), the resulting deviation from the round shape will meet the area constraint, while the perimeter of the cell increases due to the cell being stretched in the contact region (analog to surface wetting). A positive adhesion energy would conversely lead to a minimization of the contact perimeter and act like an external pressure, compressing the cell. In the interplay of different adhesion energies with different interfaces, the equilibrium cell shape results from the energetic balance of the adhesion influences and the perimeter and area constraints. Possible deviations in perimeter from the target perimeter Δ*P* = *P*− *P*_0_ serve as a measure for stretch of the cells and can be used as a biomechanical proliferation signal. Whenever Δ*P* exceeds a threshold, a proliferation step is performed, increasing the cell perimeter depending on the local tissue tension Δ*P*. To prevent unequilibrated cell shapes and uncontrolled proliferation driven by random fluctuations of the cell perimeter, we introduce a combination of parameters that define the growth function in our CPM model. The growth threshold *G*_th_ and the timescale *τ* determine how strong the mean stretching Δ*P*_*τ*_ of the cell perimeter must at least be during a time interval *τ* for growth to occur. *G*_th_ defines a relative fraction of the current target perimeter *P*_0_ and therefore the absolute threshold depends on the cell size. If the growth condition Δ*P*_*τ*_ *> G*_th_ · *P*_0_ is satisfied, the cell increases in size as follows:

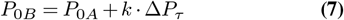

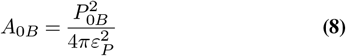

The average stretching Δ*P*_*τ*_ of the cell perimeter on the time scale *τ* is used to prevent growth based on random fluctuations. Supplementary to the averaging time scale *τ*, we also introduce the growth factor *k <* 1, which rescales the growth of each proliferation step to avoid very fast proliferation that could prevent the equilibration of cell shapes on the time scale of *τ*. The rate of the performed proliferation steps is defined by 1*/τ*. As soon as the cell has doubled in size relative to the initial size (*A*_0_ = 2*A*_0*i*_), the cell divides along its minor axis, which results in two cells with *A*_0*i*_ and *P*_0*i*_.

## Results

### Single Cell Shape

We start by simulating a single cell adhering to a substrate to determine realistic regimes for the parameters of the simulation. The influence of substrate and medium adhesion on cell shape is shown in Fig. 2. We set *T* = 1 to enable a clear understanding of other energy parameters in units of temperature. For all simulations, the elasticities for area and perimeter constraints have values of *λ*_*A*_, *λ*_*P*_ = 1, and the neighborhood order is 10. Based on eq. (6) the target perimeter *P*_0_ is defined with a static cell area of *A*_0_ = 400 *px*. The perimeter prefactor for the correction of discretiziation errors has a value of 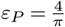 (For more detailed information on the perimeter discretization, see Supplementary Note 1). The initial condition of the simulations is a square cell with an edge length of 20 px that rests directly on a fixed substrate with a thickness of 10 px.

**Fig. 2.**
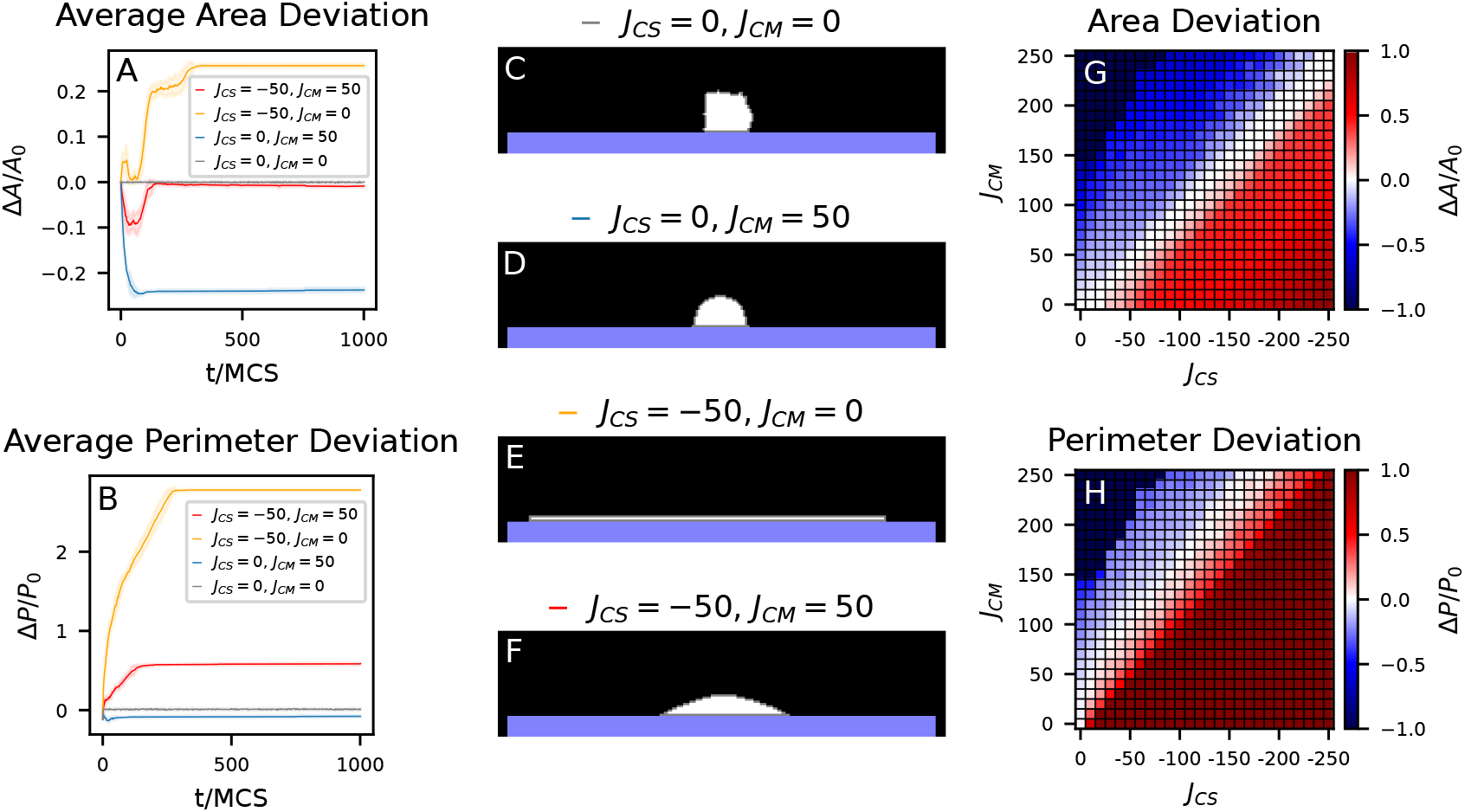
Averaged evolution of cell perimeter (A) and area (B) for different combinations of cell-substrate energy *J*_CS_ and cell-medium energy *J*_CM_ (mean and standard deviation with *N* = 10 for each condition). The equilibrium state of the simulations at MCS 1000 (C-F) shows deviations from a spherical cell shape induced by different adhesion strength. An increase in substrate adhesion leads to flattening of the cell while increasing perimeter and area. In comparison, positive cell-medium-interaction leads to a compression of the cell yielding a decreased cell area and perimeter and a minimized cell-medium-interface (D). A combination of positive *J*_CM_ and negative *J*_CS_ also results in a rounded cell-medium-interface, but the negative cell-substrate-adhesion partially counteracts the compression due to positive cell-medium-adhesion (F). G and H show the equilibrium perimeter and area for cells under the influence of different adhesion energies *J*_CS_ and *J*_CM_, showing different regimes for cell shape. In the dark blue region, where P and A are both zero, the unfavorable cell-medium-interaction dominates, leading to disappearance of the cell. In the dark red region, cell-substrate-interaction is dominant, leading to an extremely flattened cell with *P* ≥ 2*P*_0_. G also shows a white transition zone for perimeter values between these two extreme cases, where the cell perimeter fulfills the perimeter constraint *P* = *P*_0_. Mapping the transition zones between the perimeter regimes to the area plot (H) reveals that there is a region in the adhesion-parameter-space where the area of the cells fulfills the area constraint (*A* = *A*_0_), while the perimeter of the cell is increased (*P > P*_0_) without extreme flattening of the cell. This region represents an uncompressed cell on the substrate, which is stretched and can therefore grow.

Compared to an equilibrated spherical cell in the absence of adhesion (Fig. 2 C), positive cell-medium interfacial energy (membrane tension) introduces an external pressure that compresses the cell and yields a minimized cell-medium interface (Fig. 2 D). In contrast, negative cell-substrate interfacial energy (adhesion) leads to a flatted cell shape (Fig. 2 E) while increasing the cell-substrate interface and the cell perimeter. The equilibrium cell shape is determined by the energy balance between these adhesion terms and the perimeter/area constraints. A combination of positive *J*_CM_ and negative *J*_CS_ yields an intermediate cell shape between the two cases discussed above (Fig. 2 F). Cell compression due to unfavorable cell-medium interaction gets partially counteracted by the favorable cell-substrate interaction. The temporal developments of the average cell perimeter and area (Fig. 2 A-B) reveal an equilibration process on the scale of roughly 100 MCS depending on the combination of adhesion energies. One should note that the fluctuation levels of the equilibrated perimeters are lowered under the influence of positive cell-medium interaction. The time scale of the equilibration process can serve as an estimation for reasonable timescales for the introduction of a growth function that separates the time scales for cell shape equilibration and cell proliferation. These equilibration time scales may vary for the equilibration of macroscopic tissue shape in contrast to single cell shape.

We performed systematic parameter scans of perimeter and area constraints for a wide range of positive *J*_CM_ and negative *J*_CS_, which revealed different cell shape regimes (Fig. 2 G-H). For biologically realistic adherent cell shapes, we can discard the parameter ranges where cells vanish due to high *J*_CM_, or become extremely flattened due to the domination of *J*_CS_. Instead, we focus on cell shapes that fulfill the area constraint (white region in Fig. 2 H with *A* = *A*_0_) to represent the incompressibility of cells. These cells simultaneously show a significantly increased perimeter (Fig. 2 G), corresponding to increased cortical tension that is known to drive cell proliferation. In conclusion, our simulations shows contractile adherent cells with realistic shapes if the relationship between the adhesion terms is approximately *J*_CM_ = −*J*_CS_.

### Monolayer Growth

To enable the formation of a mono-layer starting from a single cell on a flat substrate, we next introduced tension-dependent proliferation (see Materials & Methods). The influence of the parameters of the growth function is shown in Fig. 3 for the time scale *τ* (A), the growth factor *k* (B) and the growth threshold *G*_th_ (C). The growth kinetics is characterized by an approximately linear increase in tissue area that stagnates at a saturation value as soon as the available substrate surface is covered by a layer of cells. The condition for cell growth is a local increase of cell perimeter (Δ*P*_*τ*_ *> G*_th_ · *P*_0_), leading to an increase in cell size proportional to *k* · Δ*P*_*τ*_ for each proliferation step. The frequency of proliferation steps is determined by the time scale *τ*. A and B show a growth rate 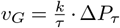 for the initial linear increase. For given interfacial energies and the resulting stretch Δ*P*_*τ*_, the fraction 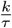 is a global scaling factor for the dynamics of proliferation relative to the equilibration of the cell shape. Similar ratios for slow growth are achieved with a large time scale *τ* or low growth factor *k*, but they result in smoother growth dynamics with low *k* and *τ* (compare, for example, the two purple curves in A and B).

**Fig. 3.**
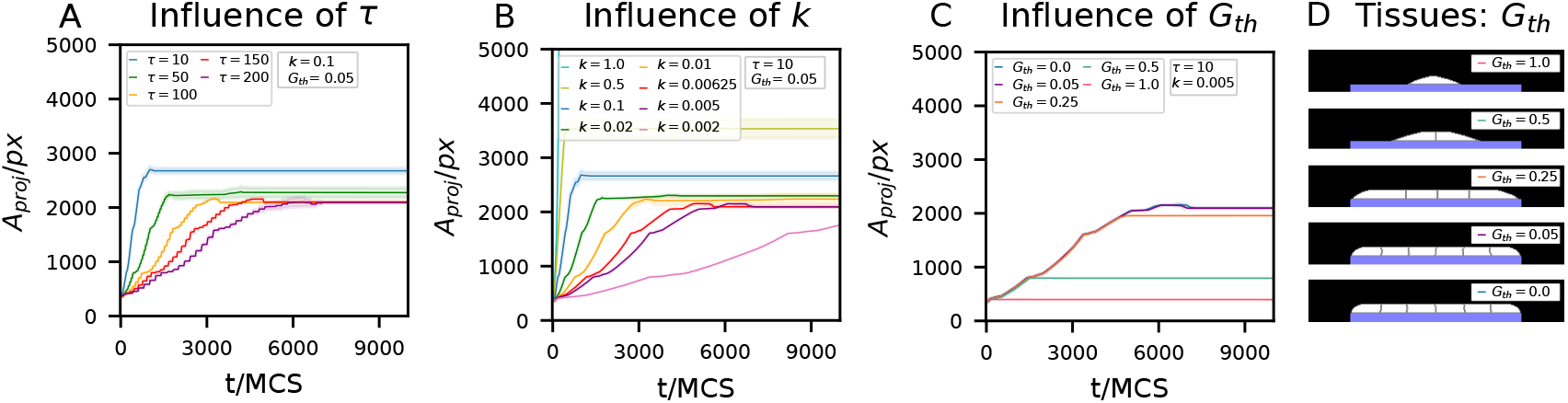
Monolayer simulations with *J*_CS_ = −50 and *J*_CM_ = 50 show the influence of the time scale *τ* (A), growth factor *k* (B), an the growth threshold *G*_th_ on the resulting proliferation dynamics. Exemplary tissues resulting from different *G*_th_ are shown next to C. The growth kinetics show an approximate linear increase in tissue area that transitions to a constant value when the available substrate is filled with cells. Subfig. A and B show the following proportionality of the growth rate: 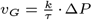.Curves with the same color in A and B have a constant value for the ratio 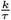 scaling the rate of growth. The comparison between A and B indicates that scaling the growth rate with *k* instead of *τ* yields smoother proliferation, which better represents continuous biological growth. Deviations from a strictly linear increase are probably related to cell divisions and a corresponding delay until the cell shape equilibrates again. The saturation values of the curves emphasize the role of a sufficiently slowly defined growth rate to separate the timescales for proliferation and shape equilibration of tissue and cells. The tissue area shows an overshoot for very fast growth rates due to unequilibrated cell shapes. Fast growth rates in B (*k* ≥ 0.01 for *τ* = 10) show larger overshoots of the final projected tissue area with the extreme case of *k* ≥ 0.1 yielding unchecked proliferation that even exceeds a monolayer configuration. If the growth rate is however suffuciently slow, the saturation value of the tissue area does not depend on the growth rate. Variable growth thresholds *G*_th_ (C) illustrate how the final equilibrium shape of the tissue is determined by the activation of the growth function if the perimeter strain exceeds the defined growth threshold. A very high growth threshold of *G*_th_ = 1 suppresses the proliferation altogether, while an intermediate value of *G*_th_ = 0.5 stops proliferation after the first cell division. For the tested values, a low threshold in the interval of *G*_th_ = [0, 0.25] enables the formation of a monolayer that fills the entire substrate. A higher or lower threshold in this interval can prevent or enable proliferation steps that are based on low perimeter strain and is therefore important for the final tissue shape.

A systematic comparison of different growth rates *v*_*G*_ reveals different saturation values for tissue area, resulting in an increased thickness of the monolayer for faster growth rates on the same available substrate area. Extremely fast growth rates (Fig. 3 B with *k* = 1) lead to uncontrolled proliferation that exceeds the monolayer configuration (data not shown). For sufficiently small growth rates, however, the saturation value remains constant. These results emphasize the importance of a suitably defined growth rate that allows for the shape equilibration of tissue and cells. Another interesting observation is that a very low growth threshold (*G*_th_ *>* 0.05) does not result in uncontrolled growth within the scope of our simulations. Note however that simulations without a threshold (*G*_th_ = 0) lead to purely fluctuation-driven, uncontrolled growth on long time scales (see supplementary Fig. S2).

Next to the suppression of purely fluctuation-driven growth, the growth threshold defines the final achievable tissue shape. Growth dynamics that rely on low Δ*P* values therefore require a low *G*_th_. Depending on the membrane fluctuation level *T* of the simulation, a suitable threshold has to be chosen to enable the desired growth dynamics while suppressing artifacts from fluctuation-driven growth. This motivates our choice to define the membrane fluctuation level *T* relatively low compared to the energy terms, which yields stiffer dynamics compared to typical CPM simulations.

### Bulk Growth

On flat substrates, tissue growth is confined to a monolayer of cells on the available area. This changes if the underlying substrate geometry allows for the formation of a concave, curved cell-medium interface. Fig. 4 A shows the resulting tissue on flat substrates on all four sides of the simulation box, with the adjustment that the substrates now touch in the lower right corner but remain spatially separated otherwise. The parameters of the growth function were the same as in Fig. 3 with *τ* = 10, *k* = 0.005, and *G*_th_ = 0.05 to separate the time scales of growth and shape equilibration. The time course of the simulation shows that, in contrast to the confined monolayer growth on the isolated flat substrates, bulk tissue now emerges from the lower right corner where the two substrates touch. The growth rate is clearly related to the local tissue curvature, as the line tension induced by the unfavorable cell-medium interaction *J*_CM_ drives the minimization of the tissue contour length. This interpretation is supported by the observation that the final tissue shape exhibits a straight contour between the anchor points of the substrate. The formation of this minimum contour length is the 2D analog to the minimal surfaces observed in 3D. Systematic variation of the adhesion energies *J*_CS_ and *J*_CM_ in Fig. 4 B and the temporal development of the tissue area in Fig. 4 C reveal how curvature-driven growth depends on the choice of interfacial energies. The parameter scan shows different types of tissue development for different parameter regimes. Cell-substrate adhesion is required for the formation of an initial monolayer. In the absence of cell-medium interaction, growth exceeding the monolayer does not occur. When both interfacial energies act simultaneously, bulk growth is enabled. The ratio between the two terms has an influence on the final tissue shape and determines whether growth follows the constraints imposed by the substrate. Most notably, a well defined tissue shape emerges only if both adhesion terms have the same magnitude. The corresponding parameter regime overlaps with the parameters used for well defined single cell shapes with *A* = *A*_0_ and *P > P*_0_ in Fig. 2.

**Fig. 4.**
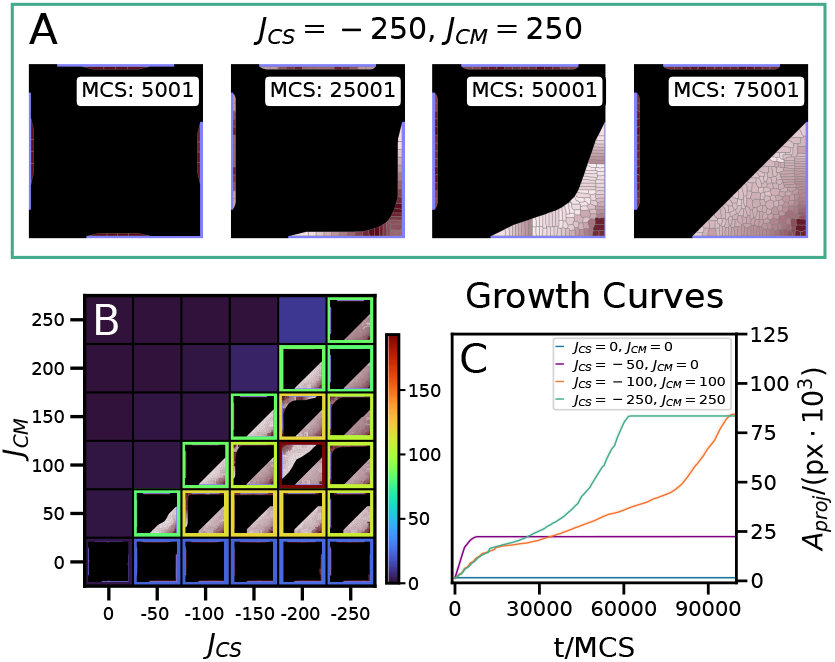
Bulk tissue growth is mediated by substrate geometry. (A) Isolated substrates on the upper and left side of the box confine growth to a monolayer, while the touching substrates of the right and bottom sides form a corner that allows bulk growth with a curved tissue-medium interface. The red colour of the individual cells depends on the relative age at the respective MCS, with a darker shade indicating older cells. (B) Tissue area and tissue shape after 100k MCS for different combinations of *J*_CS_ and *J*_CM_. This parameter scan reveals that simulations with balanced *J*_CS_ and *J*_CM_ yield bulk tissue growth in regions where the substrate is concave while being confined to a monolayer for flat substrates. Growth beyond the monolayer only occurs due to surface tension along the energetically unfavorable cell medium interface. However, an unbalanced ratio between *J*_CS_ and *J*_CM_ results in unbiological growth independent of substrate geometry. (C) compares the growth dynamics of simulations confined to a monolayer or contained bulk growth for balanced adhesion terms. In the case of balanced values for *J*_CS_ and *J*_CM_, the proliferation speed is influenced by the magnitude of both adhesion terms. The growth parameters were set to *τ* = 10, *k* = 0.005, and *G*_th_ = 0.05.

### Growth in Cleft and Capillary Geometries

To systematically investigate the influence of tissue curvature on proliferation, we analyzed the growth dynamics of clefts with various opening angles *α* and capillaries of variable width *w* (Fig. 5). The cleft geometries are inspired by the experimental setup used by Benn et al. (25). We defined different capillary geometries for the comparison of clefts to cavities with a constant width. The clefts in Fig. 5 A-C show proliferation dynamics that depend on the angle of the cleft (22.5°-90°). The data show a faster linear growth front progression for smaller cleft angles with a simultaneously larger mean curvature of the tissue contour and a higher local curvature in the center of the contour. The comparison between mean curvature and central curvature is important to verify that changes in the mean curvature are not only influenced by changes in the contour length, independent of the curvature in the center of the growth front.

**Fig. 5.**
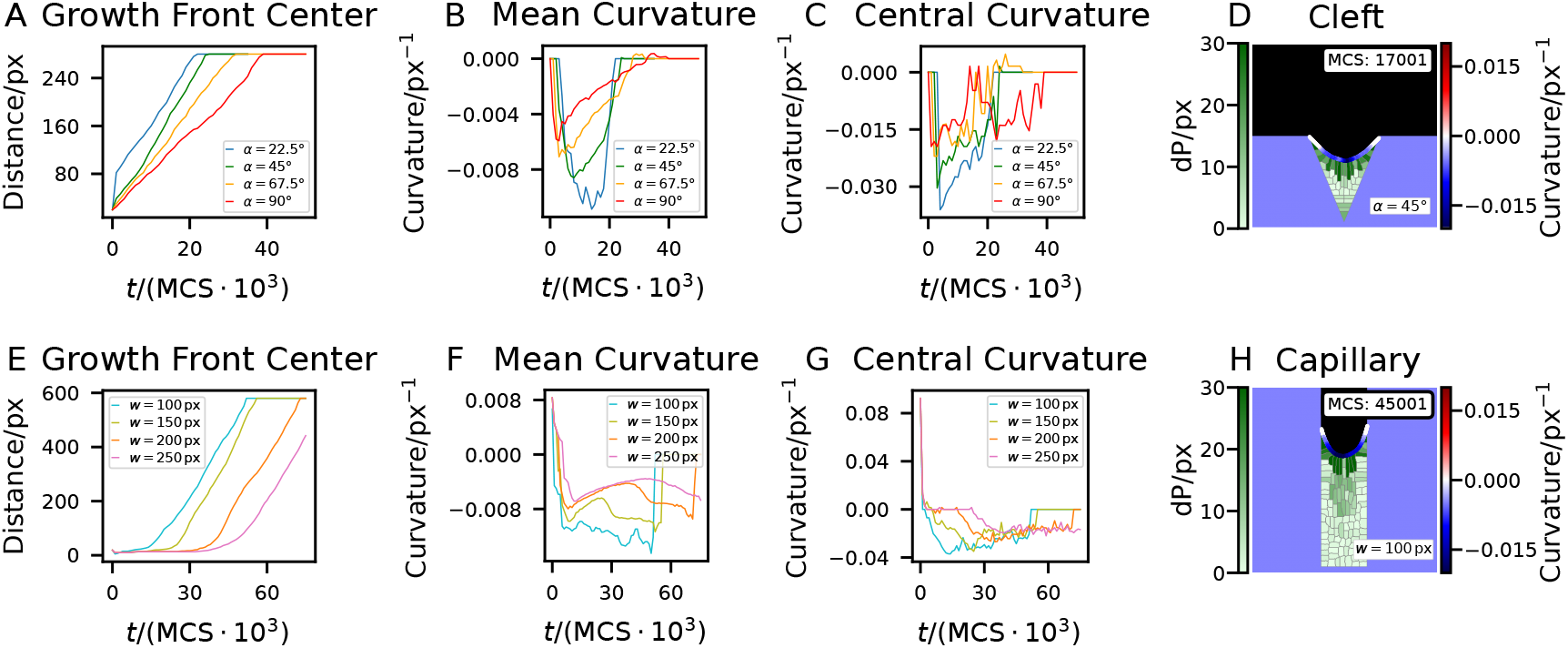
Analysis of curvature-dependent growth on various cleft and capillary substrates. Cleft geometries with different angles (A-C) reveal curvature-dependent growth dynamics. Smaller angles lead to a faster displacement of the growth front (A) and to a stronger mean curvature and center curvature (B,C). An exemplary tissue configuration in a 45°-cleft shows the color-coded local curvature at the growth front and the individual cell perimeter deviation Δ*P* in a green hue. This visualization of the local stretching of cells reveals an increased growth signal at the tissue-medium-interface. Together with the angle-dependent growth rate, this illustrates the role of local tissue curvature as a biomechanical growth signal. The curvature controlled growth of the tissue stops as soon as the contour is smoothed out. The different capillary substrates with a width between 100 px and 250 px (E-G) reveal a similar dependency between growth and curvature. The example configuration of a capillary tissue with width 150 px (H) reveals, analogous to the cleft shown, that the stretched cells that contribute to growth are localized at the curved growth front. The main difference to the growth in the cleft geometries is that, regardless of the capillary width, a constant central curvature emerges. The constant curvature that occurs after the initial transition process corresponds to a constant progression of the growth front. This supports the previous observation that growth rates depend on the local curvature. At the same time, this demonstrates the role of the underlying substrate geometry. The transition towards a constant curvature is related to the width of the cleft/capillary. The fact that the curvature between different cleft angles differs might be explained by the ongoing variation of the cleft width with the progression of the growth front. The growth parameters were set to *τ* = 10, *k* = 0.005, and *G*_th_ = 0.05 with *J*_CS_ = −250 and *J*_CM_ = 250 as adhesion parameters.

The calculation of the local curvature is based on the method used by Driscoll et al. (37) for the analysis of cell boundaries. For each tissue surface pixel of the contour, two neighbors are defined by following the contour up and down a given distance. The reciprocal radius of a circle fitted to the surface pixel and the two neighbors defines the local curvature. The result of this curvature analysis depends on the scale of the neighborhood distance and was set to 25 px to correspond to the scale of the contribution of the individual cell to the growth front. Surface pixels that are closer than 25 px to one end of the contour have a defined curvature of 0 on the given scale.

The capillary simulations (Fig. 5 E-G) show results similar to the cleft simulations with a growth front progression that is determined by the local curvature. Between the different capillary widths, differences in curvature and growth front progression occur during the initial growth phase, but then transition into a late growth phase with constant central curvature and a comparable slope in the development of growth front distance. Due to large differences in the total contour length, the mean curvature values of the different capillaries are not comparable with each other. Nevertheless, the mean curvatures also exhibit constant values for the late growth phase, corresponding to a constant progression of the growth front.

In contrast to the cleft simulations, we did not observe a difference in the late growth dynamics for different capillary widths – only the initial transition process differs in its dynamics. Based on this observation, the cleft angle dependencies can presumably be explained by the constant change in cleft width as the growth front progresses.

In summary, the cleft and capillary simulations clearly show that the local growth dynamics is determined by the local tissue curvature. The visualizations of cleft and capillary tissues in Fig. 5 D and H have a green color assigned to each individual cell indicating the strength of the local proliferation signal Δ*P*, revealing a more prevalent localization of tensed, proliferating cells at the curved cell-medium interface. These results clearly show how curvature is related to growth signaling at the single cell level and how curvature controls the resulting macroscopic tissue shape.

## Discussion

### Tension-Dependent Proliferation Reproduces Curvature-Driven Growth

Our CPM-based simulations of cells on substrates that allow the formation of a concave tissue-medium interface show bulk tissue growth for three exemplary substrate geometries. Growth on flat substrates that are spatially separated or touch in one corner of the simulation box (see Fig. 4) result in confined monolayers on isolated substrates, while bulk tissue growth emerges at concave tissue-medium interfaces. Tissue growth stops once the negative curvature of the tissue contour equilibrates to zero. The resulting minimal contour length (between the anchor points of the substrate) with zero curvature is a two dimensional analogue to a minimal surface in 3D (9, 13). Therefore, we expect the formation of minimal surfaces for CPM simulations in 3D when using a comparable tension-dependent growth function.

The simulation results of tissues growing in cleft and capillary geometries confirm these observations, and in addition imply growth rates which are proportional to the local tissue curvature (see Fig. 5). This can be explained by higher line tension in regions with higher curvature, driving proliferation to minimize the tissue contour. Differences in curvature and later growth rate are only significant between different cleft angles, but not different capillary widths. This is probably related to the instantaneous width between the anchor points of the growing tissue at a given height. In the capillary case, this width is constant, leading to constant curvature between different widths after the initial growth phase. In contrast, the width of the cleft varies dependent on height for each cleft substrate, resulting in changes in curvature and growth.

The results in Fig. 4 also reproduce the experimental observations of Kommareddy et al. on the growth dynamics of osteoblast tissue in quadratic pores (10). They found that the development of the projected tissue area is characterized by an initial lag phase followed by a universal growth rate for the bulk tissue growth. The duration of the initial lag phase is determined by substrate adhesion, while the subsequent universal growth rate is independent of the substrate, but is only determined by the cells used. Our model confirms these findings, with the difference that the initial spreading depends on the combination of *J*_CS_ and *J*_CM_, while the bulk growth is determined solely by the interfacial line tension imposed by the tissue-medium interaction *J*_CM_. Note that balanced values for *J*_CS_ and *J*_CM_ are still necessary for well defined cell shapes (*A* = *A*_0_ and *P > P*_0_, see Fig. 2) to ensure that the tissue grows under geometric confinement (see Fig. 4 for examples of unconfined proliferation).

### Cell shape

The systematic simulations of single cells (Fig. 2) under the influence of adhesion and constraints for area and perimeter illustrate the biological interpretation of the different parameters: *λ*_*A*_ describes the cell’s resistance against compression and expansion, while *λ*_*P*_ introduces a line tension along the perimeter of the cell in 2D, corresponding to a surface tension in 3D. The interfacial energies *J*_CS_ and *J*_CM_ can be interpreted as differential line tensions for contact with substrate or medium that drive changes in cell shape. Negative *J*_CS_ represents the adhesion of cells to the substrate, similar to surface wetting of a water droplet on a hydrophilic surface. Positive *J*_CM_ acts as an additional line tension for cell-medium interfaces, leading to a minimization of the interface. The resulting changes in cell shape drive proliferation based on an increased cell perimeter (Δ*P > G*_th_· *P*_0_). Therefore, *J*_CM_ is the driving factor for bulk tissue growth based on line tension at concave tissue-medium interfaces, while *J*_CS_ enables the spreading of cells on substrates.

Different cell shape regimes emerge under the counteracting influence of favorable cell-substrate and unfavorable cell-medium adhesion. We assume a cell shape on the adhesive substrate as realistic if the area constraint is fulfilled (*A* = *A*_0_) and the cell cortex is under tension (*P > P*_0_). The biological motivation for the increased perimeter is central for tension-dependent proliferation in our model, where Δ*P* = *P*− *P*_0_ serves a measure for the local mechanical tension on the level of single cells. This cortical tension is interpreted as growth signal, determining the resulting tissue shape. To keep the model as simple as possible, we selected for cell shapes that fulfill the area constraint *A* = *A*_0_, such that cells behave as incompressible droplets of liquid.

The observation that tissue growth only depends on the geometric constraints imposed by the underlying substrate geometry for balanced substrate adhesion and cortical tension highlights the importance of well-defined single cell shapes. This is evident, for example, in Fig. 4 B by confined growth in a parameter regime which simultaneously fulfills the requirements *A* = *A*_0_ and *P > P*_0_ for single cells (compare with Fig. 2). The experimentally observed orientation of surface cells perpendicular to the growth front is not reproduced in our model – this configuration would be energetically unfavorable due to the high cell-medium interfacial energy. Here, the polarization of contractile cells likely plays a role, which can be incorporated into the CPM by adding subcellular elements.

### Spreading kinetics

The spreading kinetics of the monolayer in Fig. 3 reveals a linear increase of the tissue area. Due to the constant thickness of the cell layer, the projected tissue area *A*_proj_ is proportional to the length of occupied substrate in our simulation. This can be easily transferred to experimental observations on the spreading of monolayers, in which the radius of a circular monolayer increases linearly (38). This provides evidence that the tension-dependent proliferation in the model can also reproduce the spreading dynamics of biological tissue in 3D. The analysis of the various growth function parameters shows that the Δ*P* -dependent growth rate is determined by 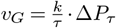,while the threshold *G*_th_ sets the minimum tension that must occur in order to activate growth. At the same time, the growth threshold prevents the occurrence of fluctuation-driven growth. The results also show that the growth rate 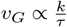 must be sufficiently slow to allow for the equilibration of cell shapes on the timescale of growth. Too fast growth leads to increased thickness of the cell monolayer in the case of spreading on a flat substrate.

Deviations of the growth kinetics from a strict linear progression are likely associated with non-linear proliferation dynamics on the single-cell level, since the enlargement of the cell depends on the relative change in perimeter, which is confined by a parabolic potential *H*_*P*_ counteracting a linear adhesion term *H*_*J*_. The systematic variations of the growth threshold suggest that the bends in the curves correspond to discrete mitosis events whose occurrence depends on the growth threshold *G*_th_. This supports the idea that deviations from a linear progression can be explained by non-linear elasticities and varying cell sizes.

It should be noted that monolayer growth on tilted substrates reveals anisotropic spreading on all substrates, that are not aligned with the lattice with selective increased growth at a substrate angle of 45°(see Fig. S3). This has implications for the development of tissues on tilted substrates, such as the clefts, shown in Fig.5 A-D. However, the main message of curvature-driven growth in the clefts should not be affected by the orientation of the substrate, as faster spreading only occurs at an angle of exactly 45°. This angle of the substrate relative to the orientation of the lattice is only present in the 90° cleft, which still shows overall slower proliferation in the cleft comparison.

### Limitations of the Model

When combining the CPM with tension-driven proliferation, some limitations need to be considered. First of all, some assumptions behind the simulations discussed here are not applicable to model more complex tissues and would instead require additional extensions and adaptations. One key point would be to transition the model from 2D to 3D. This would allow the formation of multiple directions of curvature, leading to minimal surface tissue shapes based on the geometry of the underlying substrate. In addition, the customizable properties of the CPM can be used to implement processes such as apoptosis and cell differentiation. These expansions would make the CPM more suitable to model complex tissues. Although these concepts are beyond the scope of the work presented here, the used CompuCell3D (CC3D) framework would in principle allow the implementation of such additional functionalities in an easy and user friendly manner.

Some inherent properties of the CPM can be limiting for certain applications. Those include lattice artifacts such as the anisotropy of growth on tilted substrates (Fig. S3), inconvenient parallelization (see (39) for parallelization approaches), and the lack of direct mechanical forces in the model. To rule out the possibility that the observed macroscopic growth patterns and dynamics are based on lattice artifacts, comparable simulations were rescaled at half and double resolution. Supplementary Fig. S4 shows reproducible macroscopic behavior for a wide range of lattice resolutions.

When defining CPM simulations that include tension-driven proliferation, a sensible choice of fluctuation level *T* must be made. The trade-off between simulation time and sufficiently slow growth to avoid unequilibrated cell shapes might be limiting for certain applications. Our current results imply that simulations have to be relatively stiff to prevent fluctuation-driven growth.

A possible solution to access the mechanical forces would be to calculate them from the local energy gradient defined by the energy function of the system (36). Another limitation of the CPM is that the representation of fibrous extracellular matrix (ECM) is difficult. One approach to include ECM fibers was proposed by Tsingos et al. by combining the CPM with a bead-spring model (40). This hybrid model is a promising approach to investigate the intricate feedback mechanisms between cells and ECM in the regulation of tissue homeostasis and proliferation. Overall, the tension dependent proliferation presented here could be a suitable concept to be combined with more sophisticated simulations or hybrid models to get closer to the behavior of real biological tissues.

## Conclusions

Our results highlight how a tension-dependent proliferation function implemented in the CPM framework is able to explain the macroscopic dynamics of curvature-driven tissue growth. The resulting tissue shape is determined by the external cues imposed by the geometry of the underlying substrate. Therefore, our model presents a promising approach to predict tissue shape and growth dynamics for micro-tissue engineering. The model’s customization options also allow this approach to be used to simulate individual geometries and more complex biological issues in the future. We provide predefined configurations and documentation to reproduce and adapt our simulations at https://github.com/LennardFastabend/tension-driven-cpm.

## Supplementary Note 1: Discretization of Circles

**Fig. S1.**
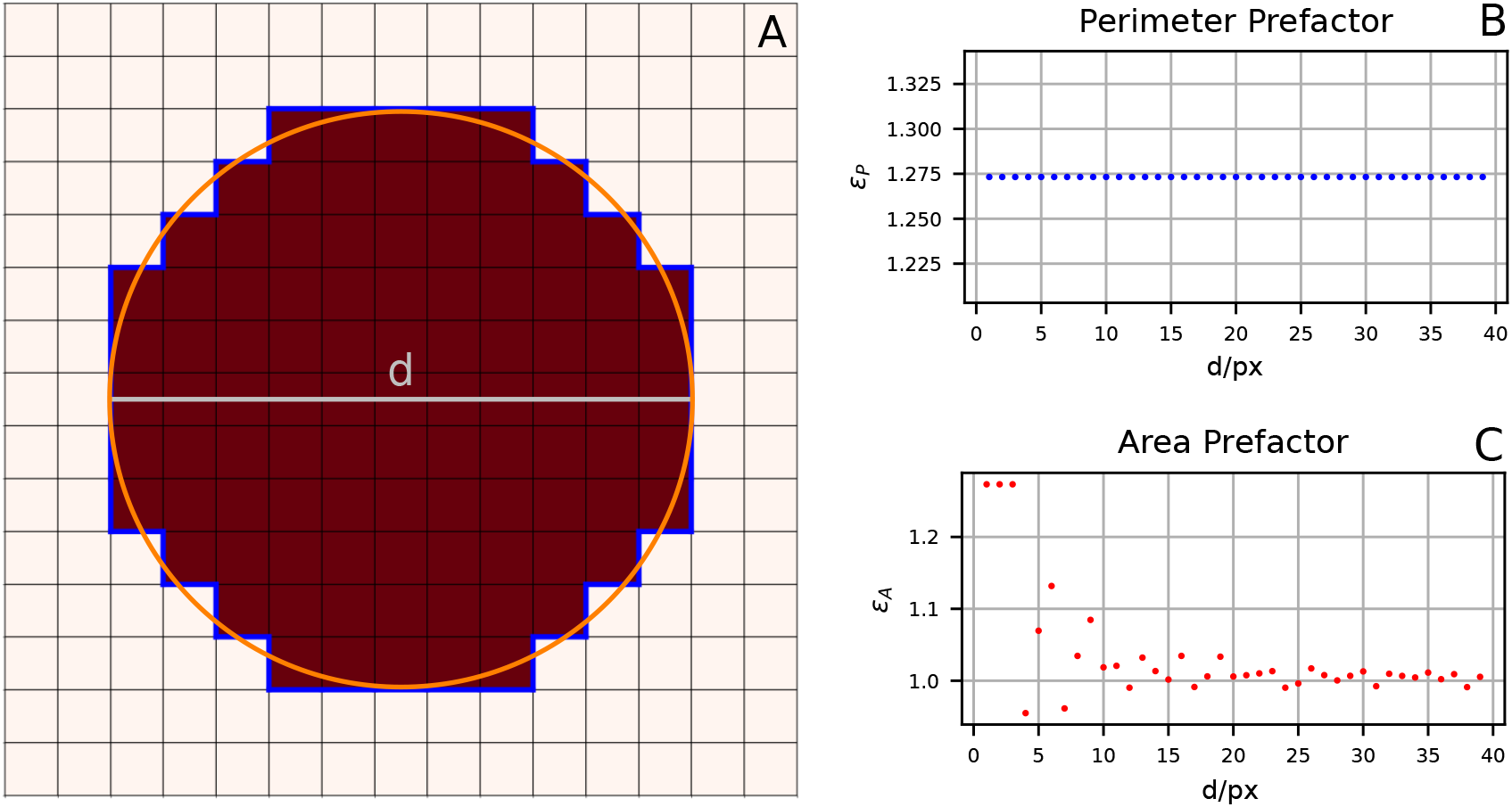
Systematic examination of the discretization error of a continuous circle onto a square lattice. A shows an example pixelated and continuous circle with a diameter of 11 px. To asses the discretization error between the continuous and pixelated perimeter, a perimeter prefactor *ε*P = *P*_*discrete*_*/P*_*continuous*_ is defined and calculated for different diameters *d*. B reveals, that the perimeter prefactor has a constant value of 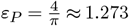 for all circle diameters. The discretization error of the area, shown in C, is defined by *ε*_*A*_ = *A*_*discrete*_*/A*_*continuous*_ and shows that the area differences for discrete and continuous circles is large for very small circles but can be neglected for larger diameters.

## Supplementary Note 2: Influence of Growth Threshold on Proliferation Dynamics

**Fig. S2.**
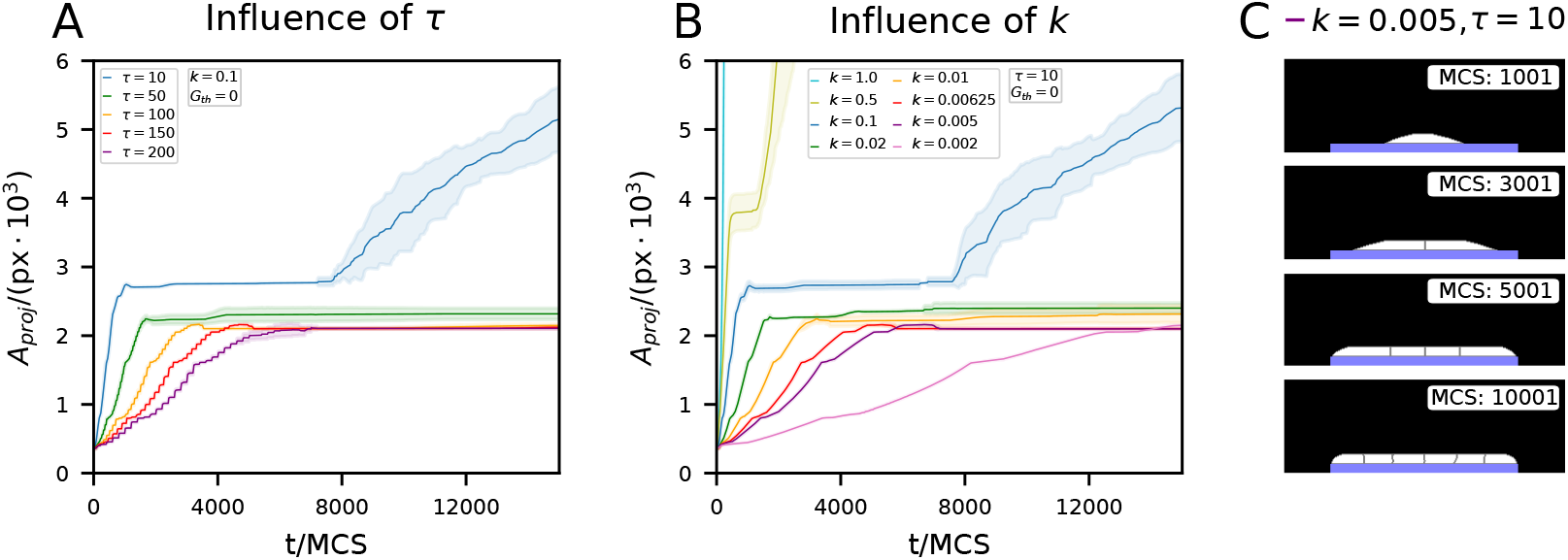
The role of the growth threshold is most clearly shown when it is omitted. The proliferation dynamics of a monolayer with *G*^th^ = 0 are shown at different growth rates by variation of time scale *τ* (A) and growth factor *k* (B) with comparable rates for similar colored curves. The adhesion parameters were set to *J*^CS^ = −50 and *J*^CM^ = 50. The development of a confined monolayer is shown exemplary in C. The growth curves reveal compared to previous results with *G*^th^ = 0.05 unconfined tissue growth after the initial saturation value is reached. B indicates that the unconfined tissue growth occurs earlier for faster growth rates. Based on these observations we decided to include a low threshold value in our model to prevent fluctuation driven growth.

## Supplementary Note 3: Anisotropy in Growth Dynamics on Serrated Edges

**Fig. S3.**
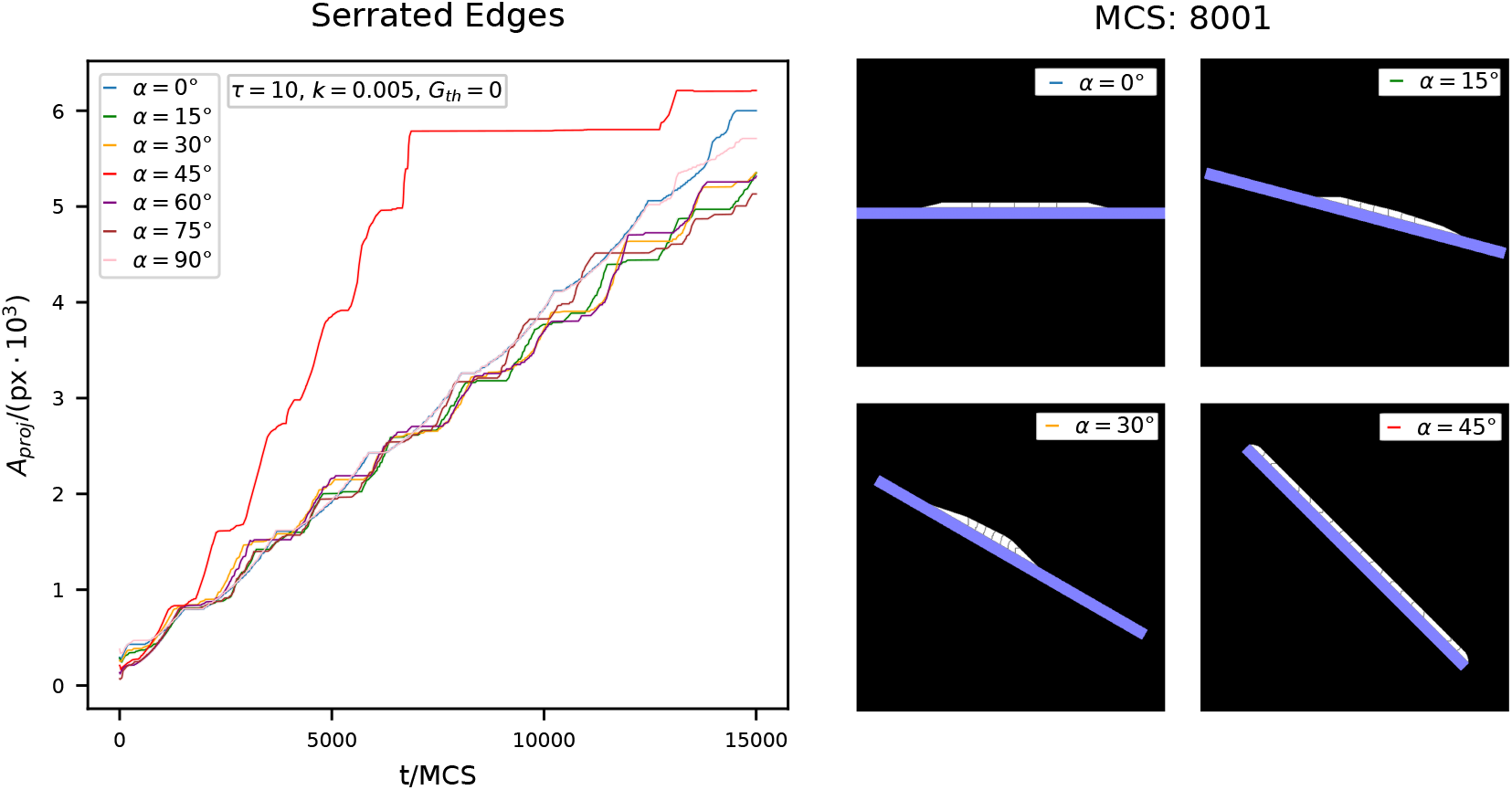
The analysis of monolayer growth on flat substrates with various tilting angles relative to the underlying lattice reveals constant projected areas of the resulting tissues with an exception for 45°. This exception is probably related to the maximized surface area of the substrate at this angle, whereas other angles, which actually also have an increased surface area do not show a difference compared to flat substrates (0°and 90°). Despite the comparable total area of the tissue cells, there are also anisotropies in monolayer growth on tilted substrates with a certain favored direction for growth. These anisotropies might influence the development of bulk tissues that rely on the development of monolayers on tilted substrates. All simulations used *J*_CS_ = −250 and *J*_CM_ = 250 as adhesion parameters.

## Supplementary Note 4: Influence of Lattice Size

**Fig. S4.**
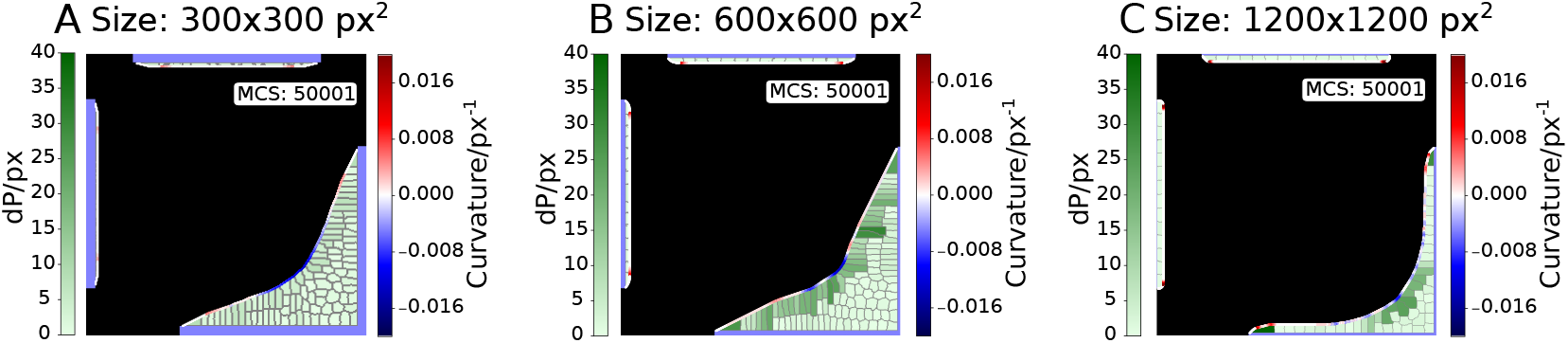
CPM simulations with different lattice sizes show comparable macroscopic growth pattern on various scales. The standard resolution (C) with a simulation box with 600 px x 600 px and an initial target cell area of 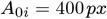 was used for all scenarios presented in this work. The halved resolution (A) with 300 px x 300 px and 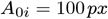 shows identical growth dynamics with a comparable tissue shape, which is merely scaled down. In contrast, the double resolution (C) with 1200 px x 1200 px and 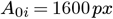 shows comparable macroscopic growth with monolayers forming on isolated substrates and bulk growth in the lower right corner of the simulation box, but the dynamic development is slower compared to the standard resolution. Despite this deviation, the comparison of different lattice sizes shows that the chosen simulation scale provides reproducible results with different sizes. It can therefore be ruled out that the tension-driven growth patterns are based solely on lattice artifacts.

## Bibliography

1. Celeste M. Nelson and Mina J. Bissell. Of Extracellular Matrix, Scaffolds, and Signaling: Tissue Architecture Regulates Development, Homeostasis, and Cancer. Annual Review of Cell and Developmental Biology, 22(Volume 22, 2006):287–309, November 2006. ISSN 1081-0706, 1530-8995. doi: 10.1146/annurev.cellbio.22.010305.104315. Publisher: Annual Reviews.

2. Jasper Foolen, Tadahiro Yamashita, and Philip Kollmannsberger. Shaping tissues by balancing active forces and geometric constraints. Journal of Physics D: Applied Physics, 49(5):053001, December 2015. ISSN 0022-3727. doi: 10.1088/0022-3727/49/5/053001. Publisher: IOP Publishing.

3. Monika Rumpler, Alexander Woesz, John W. C. Dunlop, Joost T. van Dongen, and Peter Fratzl. The effect of geometry on three-dimensional tissue growth. Journal of the Royal Society, Interface, 5(27):1173–1180, October 2008. ISSN 1742-5689. doi: 10.1098/rsif.2008.0064.

4. Pascal Joly, Georg N. Duda, Martin Schöne, Petra B. Welzel, Uwe Freudenberg, Carsten Werner, and Ansgar Petersen. Geometry-Driven Cell Organization Determines Tissue Growths in Scaffold Pores: Consequences for Fibronectin Organization. PLOS ONE, 8(9):e73545, September 2013. ISSN 1932-6203. doi: 10.1371/journal.pone.0073545. Publisher: Public Library of Science.

5. Manuela Herklotz, Marina C. Prewitz, Cécile M. Bidan, John W. C. Dunlop, Peter Fratzl, and Carsten Werner. Availability of extracellular matrix biopolymers and differentiation state of human mesenchymal stem cells determine tissue-like growth in vitro. Biomaterials, 60:121–129, August 2015. ISSN 0142-9612. doi: 10.1016/j.biomaterials.2015.04.061.

6. Cécile M. Bidan, Philip Kollmannsberger, Vanessa Gering, Sebastian Ehrig, Pascal Joly, Ansgar Petersen, Viola Vogel, Peter Fratzl, and John W. C. Dunlop. Gradual conversion of cellular stress patterns into pre-stressed matrix architecture during in vitro tissue growth. Journal of The Royal Society Interface, 13(118):20160136, May 2016. doi: 10.1098/rsif.2016.0136. Publisher: Royal Society.

7. Maike Werner, Sébastien B. G. Blanquer, Suvi P. Haimi, Gabriela Korus, John W. C. Dunlop, Georg N. Duda, Dirk W. Grijpma, and Ansgar Petersen. Surface Curvature Differentially Regulates Stem Cell Migration and Differentiation via Altered Attachment Morphology and Nuclear Deformation. Advanced Science (Weinheim, Baden-Wurttemberg, Germany), 4(2):1600347, February 2017. ISSN 2198-3844. doi: 10.1002/advs.201600347.

8. Philip Kollmannsberger, Cécile M. Bidan, John W. C. Dunlop, Peter Fratzl, and Viola Vogel. Tensile forces drive a reversible fibroblast-to-myofibroblast transition during tissue growth in engineered clefts. Science Advances, 4(1):eaao4881, 2018. doi: 10.1126/sciadv.aao4881.

9. S. Ehrig, B. Schamberger, C. M. Bidan, A. West, C. Jacobi, K. Lam, P. Kollmannsberger, A. Petersen, P. Tomancak, K. Kommareddy, F. D. Fischer, P. Fratzl, and John W. C. Dunlop. Surface tension determines tissue shape and growth kinetics. Science Advances, 5(9):eaav9394, 2019. doi: 10.1126/sciadv.aav9394.

10. Krishna P. Kommareddy, Claudia Lange, Monika Rumpler, John W. C. Dunlop, Inderchand Manjubala, Jing Cui, Karl Kratz, Andreas Lendlein, and Peter Fratzl. Two stages in three-dimensional in vitro growth of tissue generated by osteoblastlike cells. Biointerphases, 5(2):45–52, 06 2010. ISSN 1934-8630. doi: 10.1116/1.3431524.

11. C. M. Bidan, K. P. Kommareddy, M. Rumpler, P. Kollmannsberger, Y. J. M. Bréchet, P. Fratzl, and J. W. C. Dunlop. How linear tension converts to curvature: Geometric control of bone tissue growth. PLoS ONE, 7(5):e36336, 2012. doi: 10.1371/journal.pone.0036336.

12. Mario C. Benn, Simon A. Pot, Jens Moeller, Tadahiro Yamashita, Charlotte M. Fonta, Gertraud Orend, Philip Kollmannsberger, and Viola Vogel. How the mechanobiology orchestrates the iterative and reciprocal ECM-cell cross-talk that drives microtissue growth. Science Advances, 9(13):eadd9275, March 2023. doi: 10.1126/sciadv.add9275. Publisher: American Association for the Advancement of Science.

13. J. W. C. Dunlop, G. A. Zickler, R. Weinkamer, F. D. Fischer, and P. Fratzl. The Emergence of Complexity from a Simple Model for Tissue Growth. Journal of Statistical Physics, 180 (1):459–473, September 2020. ISSN 1572-9613. doi: 10.1007/s10955-019-02461-7.

14. Peter Fratzl, F Dieter Fischer, Gerald A Zickler, and John WC Dunlop. On shape forming by contractile filaments in the surface of growing tissues. PNAS Nexus, 2(1):pgac292, January 2023. ISSN 2752-6542. doi: 10.1093/pnasnexus/pgac292.

15. Andreas Roschger, Barbara Schamberger, Sebastian Ehrig, Thomas Dechat, Silvia Spitzer, Peter Fratzl, and John Dunlop. Curvature-related in-vitro formation of twisted plywood-like tissue by pre-osteoblast cells. In Osteologie, volume 32, page P 37. Georg Thieme Verlag, June 2023. doi: 10.1055/s-0043-1769682. ISSN: 1019-1291.

16. Barbara Schamberger, Sebastian Ehrig, Thomas Dechat, Silvia Spitzer, Cécile M Bidan, Peter Fratzl, John W C Dunlop, and Andreas Roschger. Twisted-plywood-like tissue formation in vitro. Does curvature do the twist? PNAS Nexus, 3(4):pgae121, April 2024. ISSN 2752-6542. doi: 10.1093/pnasnexus/pgae121.

17. Barbara Schamberger, Ricardo Ziege, Karine Anselme, Martine Ben Amar, Michał Bykowski, André P. G. Castro, Amaia Cipitria, Rhoslyn A. Coles, Rumiana Dimova, Michaela Eder, Sebastian Ehrig, Luis M. Escudero, Myfanwy E. Evans, Paulo R. Fernandes, Peter Fratzl, Liesbet Geris, Notburga Gierlinger, Edouard Hannezo, Aleš Iglič, Jacob J. K. Kirkensgaard, Philip Kollmannsberger, Lucja Kowalewska, Nicholas A. Kurniawan, Ioannis Papantoniou, Laurent Pieuchot, Tiago H. V. Pires, Lars D. Renner, Andrew O. Sageman-Furnas, Gerd E. Schröder-Turk, Anupam Sengupta, Vikas R. Sharma, Antonio Tagua, Caterina Tomba, Xavier Trepat, Sarah L. Waters, Edwina F. Yeo, Andreas Roschger, Cécile M. Bidan, and John W. C. Dunlop. Curvature in Biological Systems: Its Quantification, Emergence, and Implications across the Scales. Advanced Materials, 35(13):2206110, 2023. ISSN 1521-4095. doi: 10.1002/adma.202206110. _eprint: https://onlinelibrary.wiley.com/doi/pdf/10.1002/adma.202206110.

18. Judah Folkman and Anne Moscona. Role of cell shape in growth control. Nature, 273 (5661):345–349, June 1978. ISSN 1476-4687. doi: 10.1038/273345a0. Publisher: Nature Publishing Group.

19. Christopher S. Chen, Milan Mrksich, Sui Huang, George M. Whitesides, and Donald E. Ingber. Geometric Control of Cell Life and Death. Science, 276(5317):1425–1428, May 1997. doi: 10.1126/science.276.5317.1425. Publisher: American Association for the Advancement of Science.

20. Boris Hinz. Tissue stiffness, latent TGF-β1 Activation, and mechanical signal transduction: Implications for the pathogenesis and treatment of fibrosis. Current Rheumatology Reports, 11(2):120–126, April 2009. ISSN 1534-6307. doi: 10.1007/s11926-009-0017-1.

21. Damien Cuvelier, Manuel Théry, Yeh-Shiu Chu, Sylvie Dufour, Jean-Paul Thiéry, Michel Bornens, Pierre Nassoy, and L. Mahadevan. The universal dynamics of cell spreading. Current Biology, 17(8):694–699, 2007. ISSN 0960-9822. doi: 10.1016/j.cub.2007.02.058.

22. Thomas Lecuit and Pierre-François Lenne. Cell surface mechanics and the control of cell shape, tissue patterns and morphogenesis. Nature Reviews Molecular Cell Biology, 8(8):633–644, 2007. doi: 10.1038/nrm2222.

23. Ilka B. Bischofs, Franziska Klein, Dirk Lehnert, Martin Bastmeyer, and Ulrich S. Schwarz. Filamentous network mechanics and active contractility determine cell and tissue shape. Biophysical Journal, 95(7):3488–3496, 2008. ISSN 0006-3495. doi: 10.1529/biophysj.108.134296.

24. Wesley R. Legant, Amit Pathak, Michael T. Yang, Vikram S. Deshpande, Robert M. McMeeking, and Christopher S. Chen. Microfabricated tissue gauges to measure and manipulate forces from 3d microtissues. Proceedings of the National Academy of Sciences, 106(25):10097–10102, 2009. doi: 10.1073/pnas.0900174106.

25. Mario C. Benn, Simon A. Pot, Jens Moeller, Tadahiro Yamashita, Charlotte M. Fonta, Gertraud Orend, Philip Kollmannsberger, and Viola Vogel. How the mechanobiology orchestrates the iterative and reciprocal ecm-cell cross-talk that drives microtissue growth. Science Advances, 9(13):eadd9275, 2023. doi: 10.1126/sciadv.add9275.

26. Alexander G. Fletcher, Miriam Osterfield, Ruth E. Baker, and Stanislav Y. Shvartsman. Vertex Models of Epithelial Morphogenesis. Biophysical Journal, 106(11):2291–2304, June 2014. ISSN 0006-3495. doi: 10.1016/j.bpj.2013.11.4498.

27. Reza Farhadifar, Jens-Christian Röper, Benoit Aigouy, Suzanne Eaton, and Frank Jülicher. The influence of cell mechanics, cell-cell interactions, and proliferation on epithelial packing. Current biology: CB, 17(24):2095–2104, December 2007. ISSN 0960-9822. doi: 10.1016/j.cub.2007.11.049.

28. P. Van Liedekerke, M. M. Palm, N. Jagiella, and D. Drasdo. Simulating tissue mechanics with agent-based models: concepts, perspectives and some novel results. Computational Particle Mechanics, 2(4):401–444, December 2015. ISSN 2196-4386. doi: 10.1007/s40571-015-0082-3.

29. Tim Liebisch, Armin Drusko, Biena Mathew, Ernst H. K. Stelzer, Sabine C. Fischer, and Franziska Matthäus. Cell fate clusters in ICM organoids arise from cell fate heredity and division: a modelling approach. Scientific Reports, 10(1):22405, December 2020. ISSN 2045-2322. doi: 10.1038/s41598-020-80141-3. Publisher: Nature Publishing Group.

30. François Graner and James A. Glazier. Simulation of biological cell sorting using a two-dimensional extended potts model. Phys. Rev. Lett., 69:2013–2016, Sep 1992. doi: 10.1103/PhysRevLett.69.2013.

31. James A. Glazier, Ariel Balter, and Nikodem J. Poplawski. Magnetization to Morphogenesis: A Brief History of the Glazier-Graner-Hogeweg Model, pages 79–106. Birkhäuser Basel, Basel, 2007. ISBN 978-3-7643-8123-3. doi: 10.1007/978-3-7643-8123-3_4.

32. Jos Käfer, Takashi Hayashi, Athanasius F. M. Marée, Richard W. Carthew, and François Graner. Cell adhesion and cortex contractility determine cell patterning in the drosophila retina. Proceedings of the National Academy of Sciences, 104(47):18549–18554, 2007. doi: 10.1073/pnas.0704235104.

33. Philipp J. Albert and Ulrich S. Schwarz. Modeling cell shape and dynamics on micropatterns. Cell Adhesion & Migration, 10(5):516–528, 2016. doi: 10.1080/19336918.2016.1148864. PMID: 26838278.

34. Mark A. Wozniak and Christopher S. Chen. Mechanotransduction in development: a growing role for contractility. Nature Reviews Molecular Cell Biology, 10(1):34–43, 2009. doi: 10.1038/nrm2592.

35. D. Weaire J. Wejchert and J. P. Kermode. Monte carlo simulation of the evolution of a two-dimensional soap froth. Philosophical Magazine B, 53(1):15–24, 1986. doi: 10.1080/13642818608238968.

36. Elisabeth G. Rens and Leah Edelstein-Keshet. From energy to cellular forces in the cellular potts model: An algorithmic approach. PLoS Computational Biology, 15(12):e1007459, 2019. doi: 10.1371/journal.pcbi.1007459.

37. Meghan K. Driscoll, Colin McCann, Rael Kopace, Tess Homan, John T. Fourkas, Carole Parent, and Wolfgang Losert. Cell shape dynamics: From waves to migration. PLOS Computational Biology, 8(3):1–10, 03 2012. doi: 10.1371/journal.pcbi.1002392.

38. Antonio Brú, Juan Manuel Pastor, Isabel Fernaud, Isabel Brú, Sonia Melle, and Carolina Berenguer. Super-rough dynamics on tumor growth. Phys. Rev. Lett., 81:4008–4011, Nov 1998. doi: 10.1103/PhysRevLett.81.4008.

39. Shabaz Sultan, Sapna Devi, Scott N. Mueller, and Johannes Textor. A parallelized cellular potts model that enables simulations at tissue scale. arXiv, 2023. doi: 10.48550/arXiv.2312.09317.

40. Erika Tsingos, Bente Hilde Bakker, Koen A.E. Keijzer, Hermen Jan Hupkes, and Roeland M.H. Merks. Hybrid cellular potts and bead-spring modeling of cells in fibrous extracellular matrix. Biophysical Journal, 122(13):2609–2622, 2023. ISSN 0006-3495. doi: 10.1016/j.bpj.2023.05.013.

